# Annexin a2 as a target protein for chlorogenic acid in human lung cancer A549 cells

**DOI:** 10.1101/2020.06.11.146027

**Authors:** Lei Wang, Hongwu Du, Peng Chen

## Abstract

Chlorogenic acid, an important active component of coffee with anti-tumor activities, has been found for many years. However, the lack of understanding about its target proteins greatly limits the exploration of its anti-tumor molecular mechanism and clinical application. Here, in vitro and animal experiments showed that chlorogenic acid had a significant inhibitory effect on the proliferation of A549 cells. Using the spontaneous fluorescence characteristic of chlorogenic acid to screen the target proteins cleverly to avoid the problem of chemical modification increasing false positive, we identify and verify annexin A2 (ANXA2) as a covalent binding target of chlorogenic acid in A549 cells. Then, we discover that chlorogenic acid as an inhibitor of the binding of ANXA2 to p50 subunit inhibited the expression of downstream anti-apoptotic genes cIAP1 and cIAP2 of NF-κB signaling pathway in A549 cells in vitro and vivo. Moreover, we find chlorogenic acid hindered the binding of ANXA2 and actin maybe involved in the impediment of tumor cell cycle and migration. Thus, this work demonstrates that chlorogenic acid, as a binding ligand of ANXA2, decrease the expression of NF-κB downstream anti-apoptotic genes, inhibiting the proliferation of A549 cells in vivo and vitro.

## Introduction

Natural products provide abundant and effective small molecule drugs for the treatment of human diseases. As one of natural products, chlorogenic acid (CGA), a phenolic acid synthesized from caffeic acid and quinine acid, is widely found in people’s diet, cosmetics and medicines (1). Actually, CGA is considered to be one of the active substances of many traditional Chinese medicines such as *Eucommia ulmoides, honeysuckle* and mung coffee bean (2) with outstanding efficacy in anti-tumor, anti-bacterial and anti-inflammatory.

In the early stage, the pharmacological value of CGA was found to be antiviral and anti-inflammatory. CGA has been proved plays a significant role in inhibiting bacteria and viruses (3,4), alleviating and eliminating inflammation (5) in vitro and in vivo. CGA and its derivatives show inhibition in the fight against hepatitis B and hepatitis C (6,7). After a thorough study of the antimicrobial, antiviral and anti-inflammatory effects of CGA, it has been proved that some important signaling pathways involved in these processes (8–10). Among them, the value of NF-kB pathway widely recognized. Most remarkable, in the field of cancer which seriously endangers people’s health, there are many experimental evidences that CGA has a significant inhibitory effect on cancer. The anti-tumor test of chlorogenic acid has entered the clinical trial stage in China. The results of phase I/II clinical studies show that it has great safety and tolerance in clinical trials, and there is no obvious side effect. These advances show that CGA has a greater application prospect in the treatment of tumors.

Lung cancer is one kind of the highest incidence and mortality rate cancer in China. New drug demand for lung cancer treatment in is urgent (11). The consensus that CGA can inhibit the proliferation and invasion of human lung cancer A549 cells and other cancer cells has been gradually formatted (12–14). Phenotype-based pathway research is the current research mainstream on CGA. In details, some researchers reported that CGA may inhibit the development of cancer cells through such signaling pathways, especially NF-κB (12,15,16). Some studies also found that CGA has a ignificant effect on cell cycle, it have been demonstrated that CGA can inhibit glioblastoma by transforming macrophages from M2 phenotype to M1 phenotype (17). Many previous anti-tumor studies have reported phenotype-based signaling pathways and even clinical studies of CGA, However, there are few reports about its anti-tumor mechanism based on the study of chlorogenic acid target proteins. Additionally, the researches on CGA target proteins are very limited (18–20), and almost no important target proteins related to its medicinal functions have been reported in detail.

In order to understand the anti-tumor activities of CGA more comprehensively, it is necessary to screen the target proteins directly binding to CGA. Because chlorogenic acid has a significant inhibitory effect on human lung cancer cell line A549, it was selected as a typical tumor cell model in this study. Based on the characteristics of CGA spontaneous fluorescence, we used a comprehensive and reliable screening method to identify and verification the molecular target of CGA. We hope to provide an accurate molecular mechanism for the anti-tumor activity of CGA in A549 cells.

## Results

### The effects of CGA on activities of A549 cells

According to the design of this study (Figure 1A), in order to explore the effects of CGA on activities of A549 cells, we tested the changes of proliferation of A549 cells after CGA stimulation. And we also carried out quantitative proteomics to recognize this influence. First, we discovered that the cell viability was gradually inhibited with the increase of CGA concentration according to the results of cytotoxicity test of CGA on A549 (Figure 1B).

**Figure 1.**
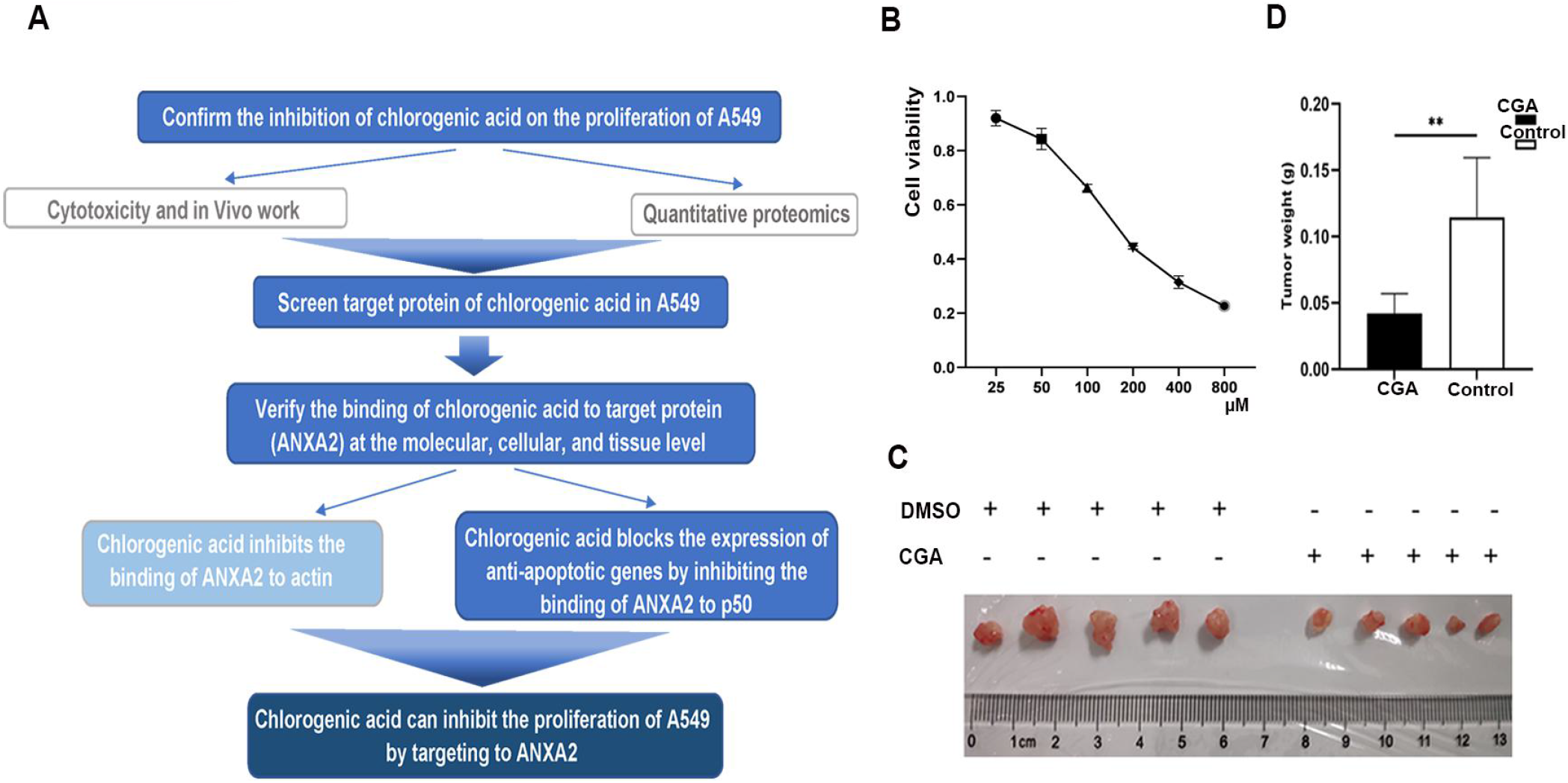
The effects of CGA on activities of A549 cells. **A.** The overall technical route of this study. **B.** The inhibitory effect of CGA on A549 cell was dose-dependent. When the CGA concentration was about 150 μM, the cell viability remained about 50%. Each experimental set is representative of n=5 and error bars as mean±SD. **C.** The tumor of the CGA treatment group and the control group were compared and photographed. CGA treatment group was obviously less than the control group (n=5). **D.** Statistical chart of tumor weight.

After CGA was proved to be effective in vitro, in vivo experiments were also carried out. Nude mice were subcutaneous injected of A549 to make subcutaneous tumor model and then continuous injected of CGA or DMSO (control) for four weeks. It was found that the tumors in CGA treatment group were significantly smaller than control group (Figure. 1C), and the mass was about half of that in the control group (Figure. 1D).

### Screening of target proteins based on the spontaneous fluorescence of CGA

Based on the above findings, we hoped to understand the mechanism through the target protein interacting with CGA. Firstly, the green fluorescence induced by the chemical structure of CGA (Figure 2A) was confirmed. After scanning by the green channel (532 nm), the CGA array showed obvious green fluorescence (Figure 2B). The green fluorescence (Figure. 2C) observed in A549 cells indicated that target proteins of CGA exist in this cell line. This is also an important reason for A549 cells was selected as the research object of this study.

**Figure 2.**
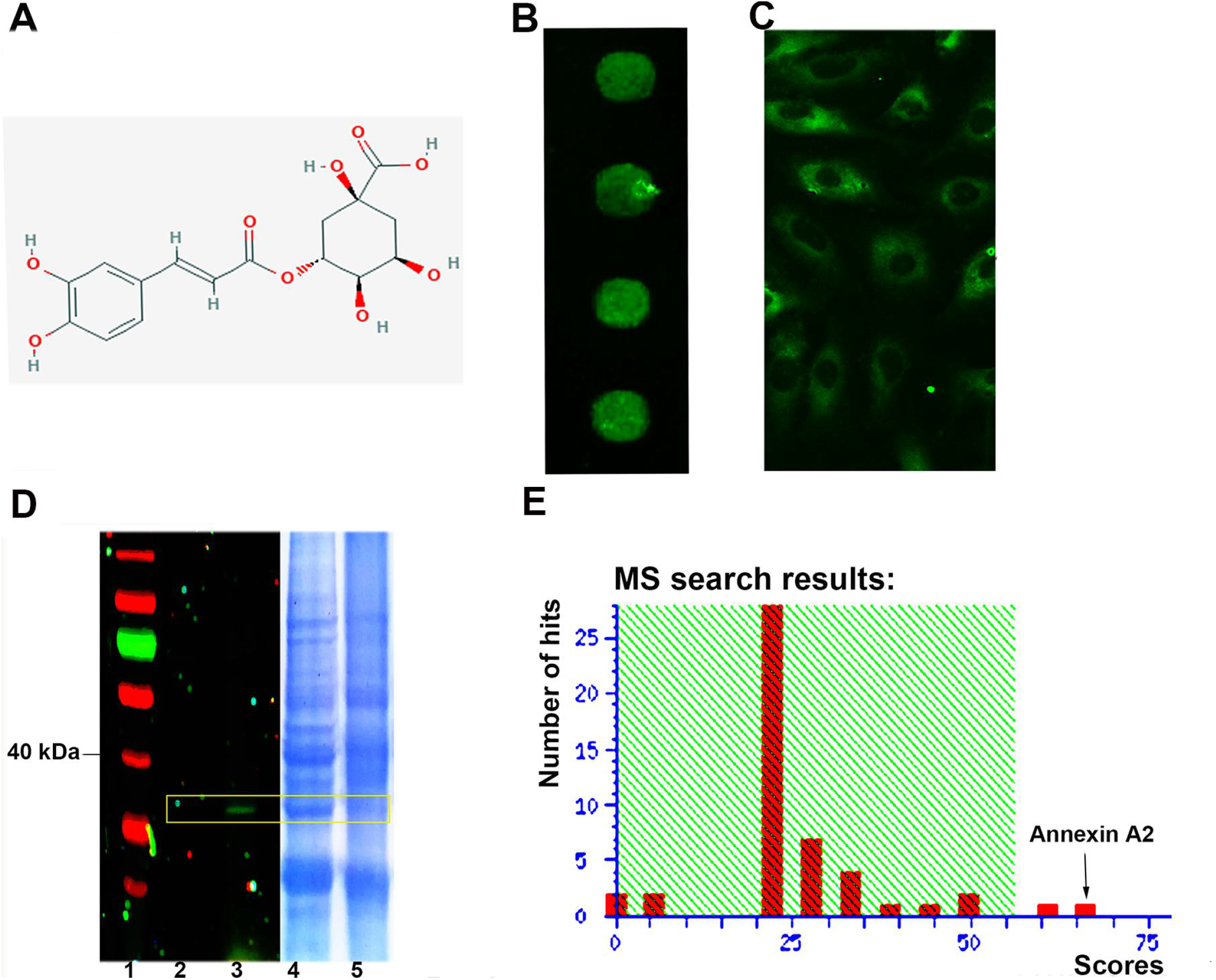
ANXA2 was identified as a target protein of CGA in A549 cells. **A.** The chemical structure of chlorogenic acid. **B.** CGA was dotted on the slide and then scanned. The brightly green fluorescence was observed. **C.** After co-incubation of CGA with A549, CGA (Green) were observed under fluorescence microscopy (20 x). **D.** The fluorescence scanning result of SDS-PAGE was imagined (Lane 1: protein ladder; Lane 2: A549 whole protein-DMSO incubation group; Lane 3: A549 whole protein-CGA incubation group; Lane 4, lane 5 were the results of Lane 2, lane 3 stained with Coomassie brilliant blue dye.). **E.** The fluorescent protein band was identified as ANXA2 by MALDI-TOF/TOF.

Given the strong fluorescence of CGA, we performed a series of screening experiments using CGA as a fluorescent probe in A549 cell lysates to identify its target proteins. As expected, the proteins with green fluorescence was found on the gel. To identify these proteins that were captured by the CGA, the fluorescent band was then cut down and digested with trypsin. The obtained peptide fragments were identified by MALDI TOF/TOF and analyzed (Figure. 2D). The protein with the highest coverage of peptide segments was considered as the potential target protein. The peptide segments of the suspected target protein (about 36 kDa) shared top homologous sequences with ANXA2 and *P*<0.05 (Figure. 2E, mass spectrogram in Figure. S1).

The phenotype based quantitative proteomics to study the influence of chlorogenic acid (CGA) on cancer cells can accurately and comprehensively represent the operation of different vital activities through the changes of different functional proteins and point out the direction for future research. Thus, we employed quantitative proteomics to recognize the influence of CGA on vital activities of tumor cells which was implemented according to the flow chart (Figure S2). More than 3100 proteins were identified by Tandem-Mass-Tag (TMT) quantitative proteomics, and about 3000 of them were quantified. The down regulated proteins included many important proteins involved in many biological processes, such as anti-apoptotic, this is consistent with the function of ANXA2 in tumor cells, so it was selected as the potential research target.

### ANXA2 is verified as a target protein of CGA at molecular level

Aimed at characterizing the CGA-ANXA2 complex in vitro, the recombinant ANXA2 was successfully expressed in *E. coli* and purified (Figure 3A, Verification results of mass spectrometer was showed in Figure S3). Afterwards, the recombinant ANXA2 with different treatment was incubated with CGA for 2h. After SDS-PAGE, the recombinant ANXA2 showed brightly green fluorescence (lane 10 and 11 in Figure 3B). Normally, BSA as control did not show any fluorescence (lane 5, 6 in Figure 3B). The ANXA2 pre-treated by high temperature had only weak fluorescence (lane 4 in Figure 3B). The ANXA2 stored at ultra-low temperature showed almost no fluorescence (lane 3 in Figure 3B). However, after urea dissolution and purification, the renatured ANXA2 also did not have any green fluorescence (lane 7 and 8 in Figure 3B). The results indicated that the binding of CGA to ANXA2 is a binding closely related to protein structure.

**Figure 3.**
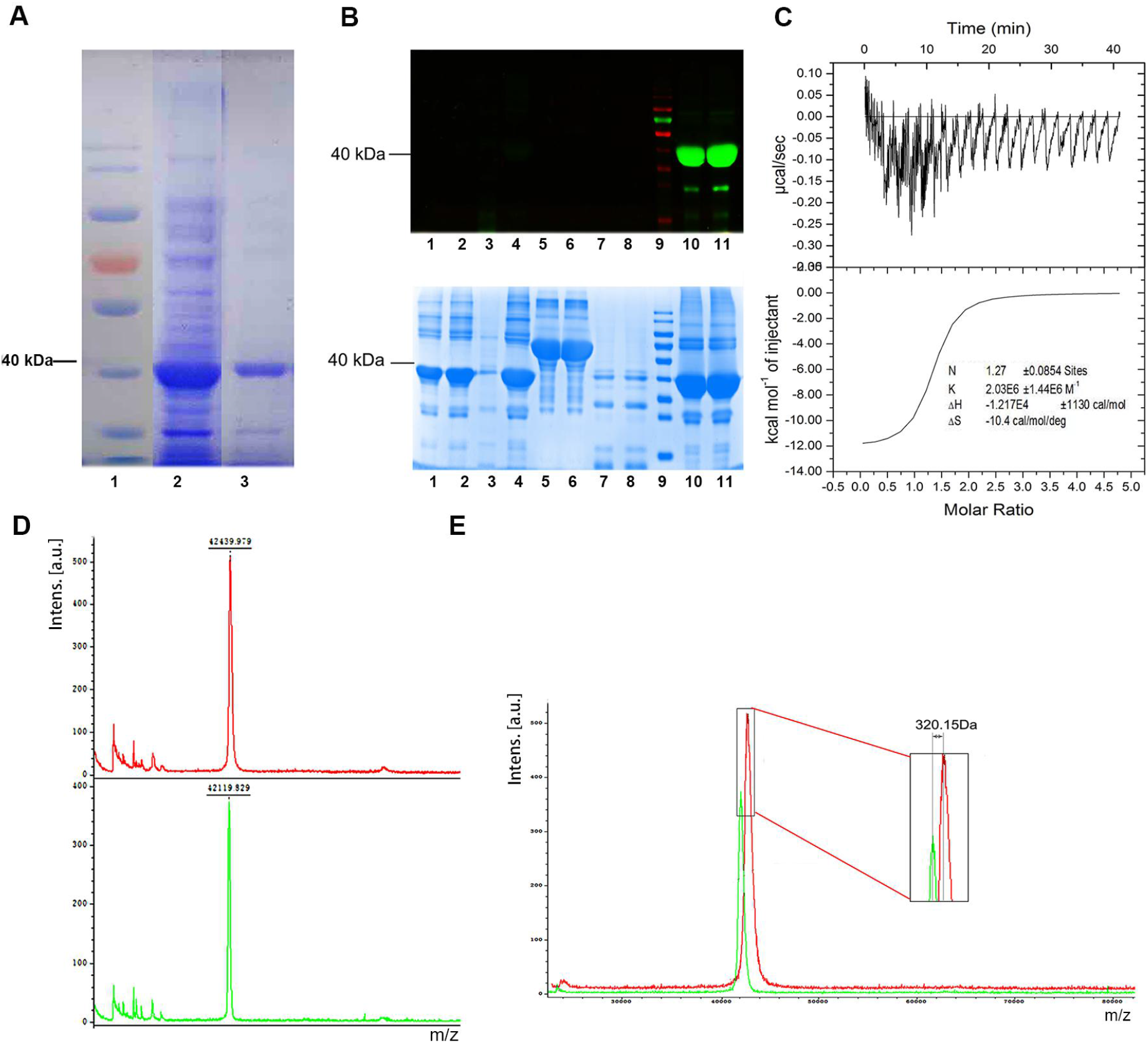
Verification of ANXA2 binding to CGA at molecular level. **A.** Recombinant ANXA2 was obtained by *E. coli* expression system. The lanes (from left to right) were protein ladder (Lane 1), whole protein of *E. coli* BL21 induced by IPTG (Lane 2) and purified ANXA2 (Lane 3). **B.** CGA was incubated with recombinant ANXA2 as described and SDS-PAGE was performed. The above picture in (**B**) is a gel graph under the fluorescence scanner; the following in (**B**) is a gel graph after Coomassie brilliant blue staining. Lane 1, 2: recombinant ANXA2; Lane 3: ANXA2 which was stored at ultra-low temperature then re-dissolution; Lane 4: The recombinant ANXA2 by heated; Lane 5, 6: BSA; Lane 7, 8: the recombinant ANXA2 with urea denaturation; Lane 9: protein ladder; Lane 10, 11: the ANXA2. **C.** Isothermal Titration Calorimetry (ITC) proved that ANXA2 binds strongly with CGA at the thermodynamic level. **D.** The molecular weights of ANXA2 (Green curve, 42119.829 Da) and ANXA2-CGA complex (Red curve, 42439.979 Da) were determined by MALDI-TOF/TOF. **E.** Comparing the two measured molecular weights, the fact that ANXA2 increased 320.15 Da after co-incubation with CGA was found.

To detect the affinity of ANXA2 and CGA, a qualitative method called Isothermal Titration Calorimetry (ITC) was used. The results showed that the binding affinity of the two molecules was relatively strong. The binding affinity and binding stoichiometry of ANXA2 to CGA can be obtained by single site binding model fitting of the integrated binding heat (Figure 3C). According to ITC, the binding affinity of a ligand can be classified into three levels: low binding affinity (Ka <10^4^M^−1^), moderate affinity (Ka = 10^4^−10^8^M^−1^), and high affinity (Ka > 10^8^M^−1^). The Ka of the two molecules was 2.03E6+1.44E6 M^−1^. This indicates that ANXA2 has a strong affinity with CGA. According to the analysis of ITC results, there may be only one site in the binding of ANXA2 to CGA (Figure 3C).

In addition, the most direct difference of ANXA2 binding to CGA is the increase of molecular weight. Therefore, after using MALDI TOF/TOF to identify and compare ANXA2 and ANXA2-CGA complex (Figure 3D), it was found that the molecular weight of the complex (Red curve) increased obviously (the two peaks were obviously dislocated). The peak representing the complex lagged behind ANXA2 (Green curve) (Figure 3E), indicating that the molecular weight of the complex was increased. The molecular weight of the complex was about 42439.979 Da, while that of ANXA2 was 42119.829 Da. The difference (320.15Da) of two molecular weights similar to that of one CGA molecule (354.31Da). This implied that ANXA2 binds to CGA in a ratio of 1:1. This result was the same as that of ITC. Comprehensive analysis, the results indicated that the binding of CGA to ANXA2 with ratio 1:1 is a covalent binding closely related to protein structure.

### Verification of ANXA2 binding to CGA at cellular and tissue level and binding site prediction

To verify complexation between CGA and ANXA2 in living cells, we treated A549 cells with CGA and then probed for endogenous ANXA2 using an immunofluorescence assay so that the localization of ANXA2 and CGA could be monitored under fluorescence microscopy. Among them, CGA showed green fluorescence and goat anti-mouse antibody labeled with Cy5 fluorescence was used to characterize the location of ANXA2 (red). The cytoplasm and cell membrane of A549 showed obvious green and red fluorescence (Figure 4A-B). Merged color (orange) showed that their position was almost the same. And the results of co-localization of the two in human lung cancer tissues also showed that they existed with identical location (Figure S4). The results of co-localization of CGA and ANXA2 at cellular and tissue levels directly verify the binding of CGA to annexin A2. Moreover, CGA can strongly bind to the different position of ANXA2 in the A549 cells.

**Figure 4.**
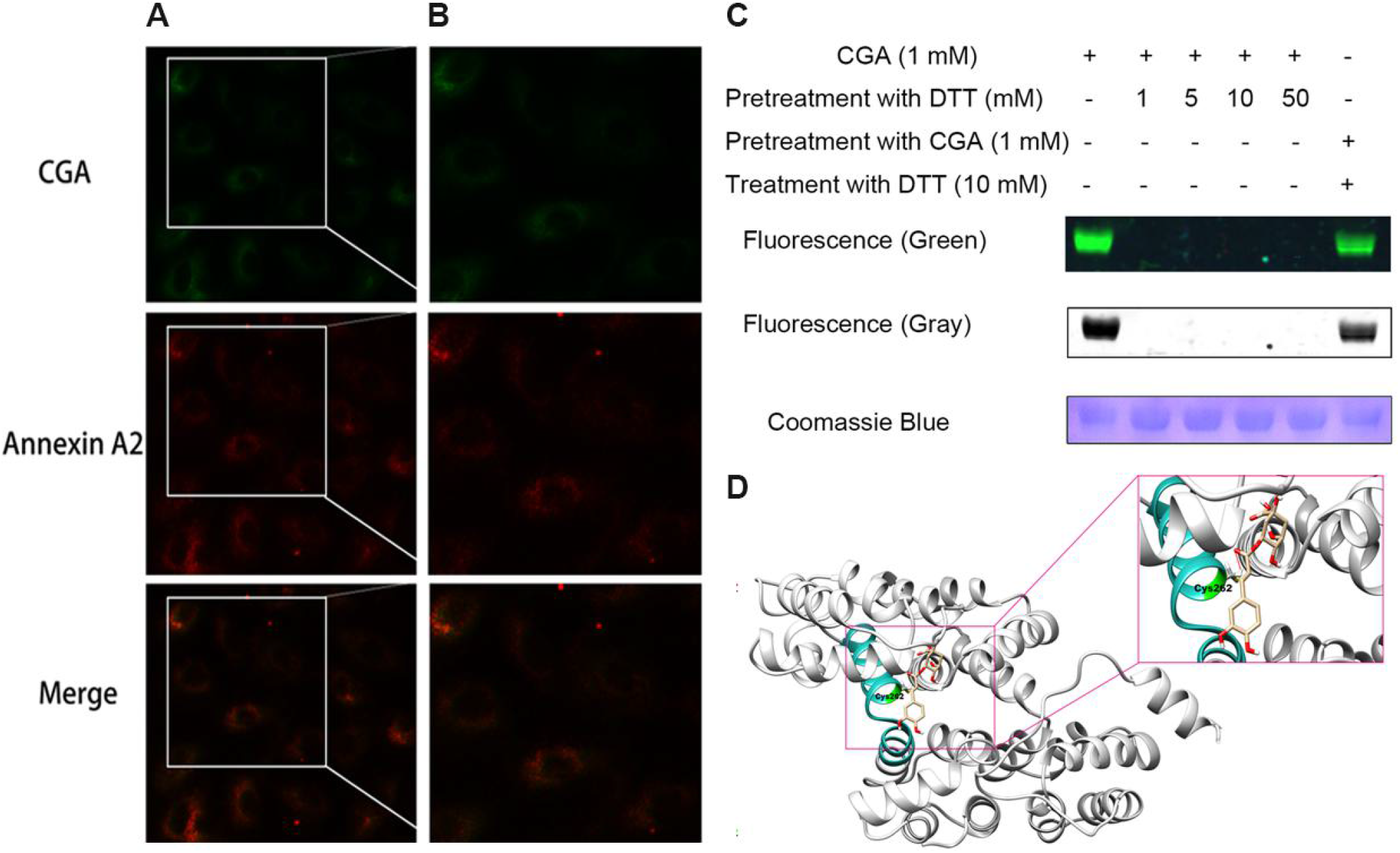
Cell level binding verification and site analysis of ANXA2 to CGA. **A.** Green fluorescence of CGA and red fluorescence of ANXA2 were observed in cell wall, cytoplasm and nucleus. The fluorescence positions of the two were basically coincident (10×). **B.** cells in (**A**) after enlargement. **C**. DTT inhibited the combination of ANXA2 with CGA. After being treated with DTT of different concentrations, ANXA2 could not combine with CGA. However, DTT treatment had no effect on ANXA2 pre-incubated with CGA. **D.** UCSF Chimera simulation results showed that CGA may bind to ANXA2 in a cavity of the protein. The covalent binding of the two might be related to cys262.

Because CGA has the structure of unsaturated ketone, we considered whether the binding is related to a similar Michael addition reaction mode that has been reported to be related to cysteine(21). When ANXA2 was pretreated with dithiothreitol (DTT), it was found that the binding efficiency of ANXA2 with CGA was inhibited, which was directly reflected in the fact that no green fluorescence was detected after incubation with CGA (Figure 4C). However, ANXA2 pre-incubated with CGA was not affected by DTT (Figure 4C). Therefore, we speculated that the covalent binding of CGA and ANXA2 was related to cysteine.

Here, we used UCSF Chimera software to simulate the binding position of CGA and ANXA2, and provided guidance for the real binding position of them. Based on the results of molecular docking simulation and experimental facts, it was inferred that CGA binds closely to annexin A2 in a cavity formed by several peptide segments. Moreover, there was the 262^nd^ cysteine (Cys262) of ANXA2 in the vicinity of CGA (Figure 4D). We speculated CGA might bound to ANXA2 covalently by cys262.

### CGA prevented ANXA2 from binding to NF-κB p50

As known, the complex formed by ANXA2 and p50 can promote the transcriptional activity of NF-κB and up-regulate the transcription of downstream target genes of NF-κB by transferring to the nucleus, thus enhancing the anti-apoptotic ability of cancer cell(22) (Figure 5A). The protein A/G beads with anti-ANXA2 antibody was used to pull down the p50 combined with ANXA2 to characterize the amount of ANXA2-p50 complex. When CGA was added to the mixed system of ANXA2-p50, the formation of ANXA2-p50 complex decreased significantly (Figure 5B). ANXA2-p50 complex in A 549 cells was also detected by Western blot between CGA stimulation and normal group. After simulated, ANXA2-p50 complex in cells was also decreased significantly (Figure 5C). The results of pull-down experiment and Western bolt showed that CGA inhibited the binding of ANXA2 to p50. Therefore CGA might more likely have a significant regulatory effect on its downstream pathways.

**Figure 5.**
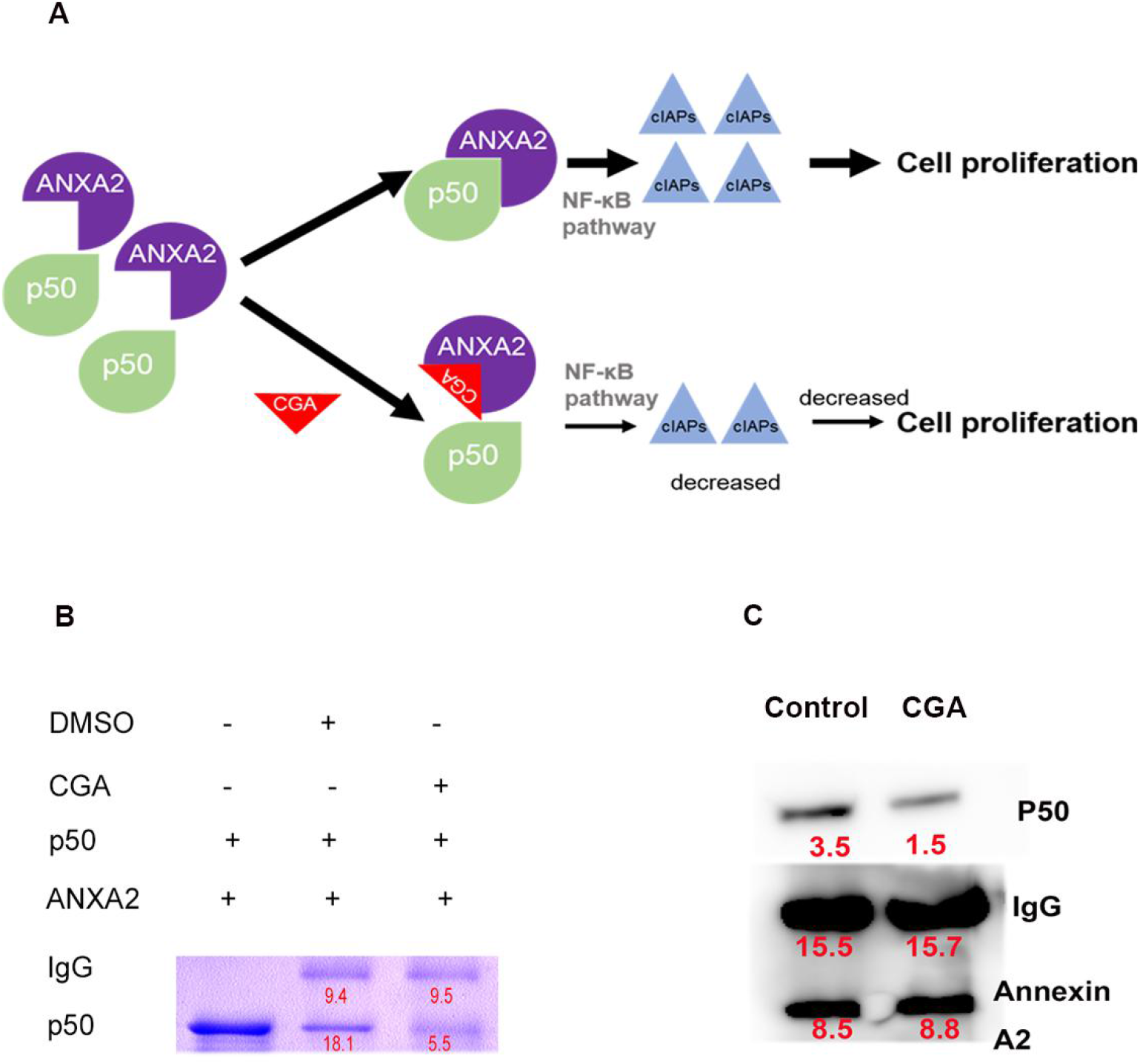
CGA prevented ANXA2 from binding to NF-κB p50 in A549 cells. **A.** CGA binding Annexin A2 may block AXNA2-p50 pathway. **B.** The inhibition of CGA on the formation of ANXA2-p50 complex was proved by co-immunoprecipitation. The addition of CGA hindered the combination of them. **C.** Stimulated by CGA, the ANXA2-p50 complexes were decreased significantly in A549 cells by co-immunoprecipitation with ANXA2 antibody, then detected by Western blot with p50 antibody.

### CGA down-regulated the expression of anti-apoptotic genes in downstream of NF-κB pathway in vitro and in vivo

After confirming that CGA blocked the binding of ANXA2 to p50, the expression of downstream anti-apoptotic genes of NF-κB pathway was detected. When A549 cells was treated with 400 μM CGA, the expressions of anti-apoptotic genes - cIAP1 & cIAP2 displayed obvious down-regulation at the mRNA level (Figure 6A). Western Blot showed that the expression of ANXA2 and p50 did not alter with CGA stimulation (Figure 6 B-C). This suggests that CGA does not play a role by directly affecting the expression of ANXA2 or p50. However, the anti-apoptotic genes-cIAP1 & cIAP2 decreased significantly (Figure 6 D-E). This suggested that CGA down-regulated the downstream anti-apoptotic genes by blocking the binding of ANXA2 to p50 in A549.

**Figure 6.**
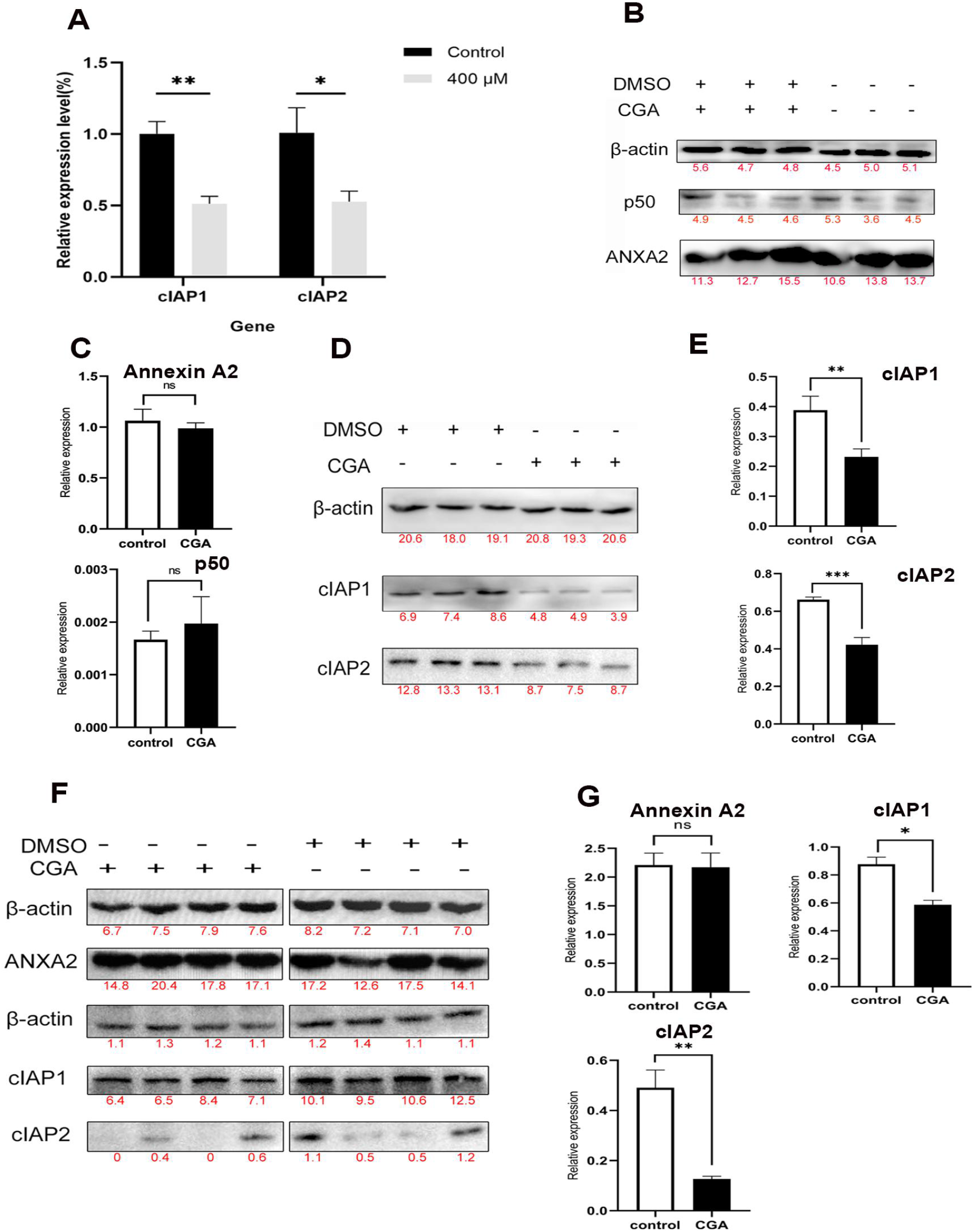
CGA inhibited the binding of ANXA2 to p50 and decreased related anti-apoptotic genes in vitro and vivo. **A.** The mRNA expression of NF-κB downstream anti-apoptosis related genes in A549 cells were significantly reduced by CGA stimulation. Each experimental set is representative of n=3 samples and error bars as mean±SD. **B.** There was no significant change in the expression of ANXA2 and p50 at protein level before and after CGA stimulation. **C.** Quantitative statistics of result in (B). **D.** After CGA stimulation in A549 cells, the expression of anti-apoptosis genes cIAP1 and cIAP2 decreased significantly at protein level. Each experimental set is representative of n=3 samples and error bars as mean ± SD. **E.** Quantitative statistics of result in (D). **F.** Western blot was used to detect the expression of related genes in mouse A549 tumor tissues including the treatment group and the control group. Each group involved four different tumor tissues (n=4). The anti-apoptotic genes cIAP1 and cIAP2 were significantly down regulated in treatment group. **G.** Quantitative statistics of result in F. Data are mean ± SEM. \**P* < 0.05, \*\**P* < 0.01, \*\*\**P* < 0.001.

In order to explore whether the anti-apoptotic genes down-regulated by NF-kB pathway was same as the results in vitro, the expressions of cIAP1 and cIAP2 of tumor tissue in animal work were detected by Western Blot. The results illustrated that the two anti-apoptotic genes were significantly down regulated (Figure 6 F-G). This was consistent with results in vitro. This emphasized that CGA can inhibit tumor development by down regulating the anti-apoptotic genes in downstream of NF-κB pathway in vivo.

### CGA blocked the binding of ANXA2 to actin involved in cell cycle

We design a co-immunoprecipitation (Co-IP) experiment to find the effect of CGA on other proteins except NF-κ B p50 that interact with ANXA2(Figure 7A). After SDS-PAGE of proteins pulled down by protein A/G, the results showed that the number of protein bands in the control group was more than that in the CGA pretreatment group (Figure 7B). The results of the two methods (intracellular and extracellular) were the same. The band was identified by LC-MS / MS. The protein band more than 40 kDa was identified as actin (top Sequence coverage 26.7% with 9 special peptides). Blocking their binding by CGA might inhibit the cell cycle and migration of tumor cells (Figure 7C). Because actin is important skeleton protein, which play an important role in the formation, division, and movement of cancer cell.

**Figure 7.**
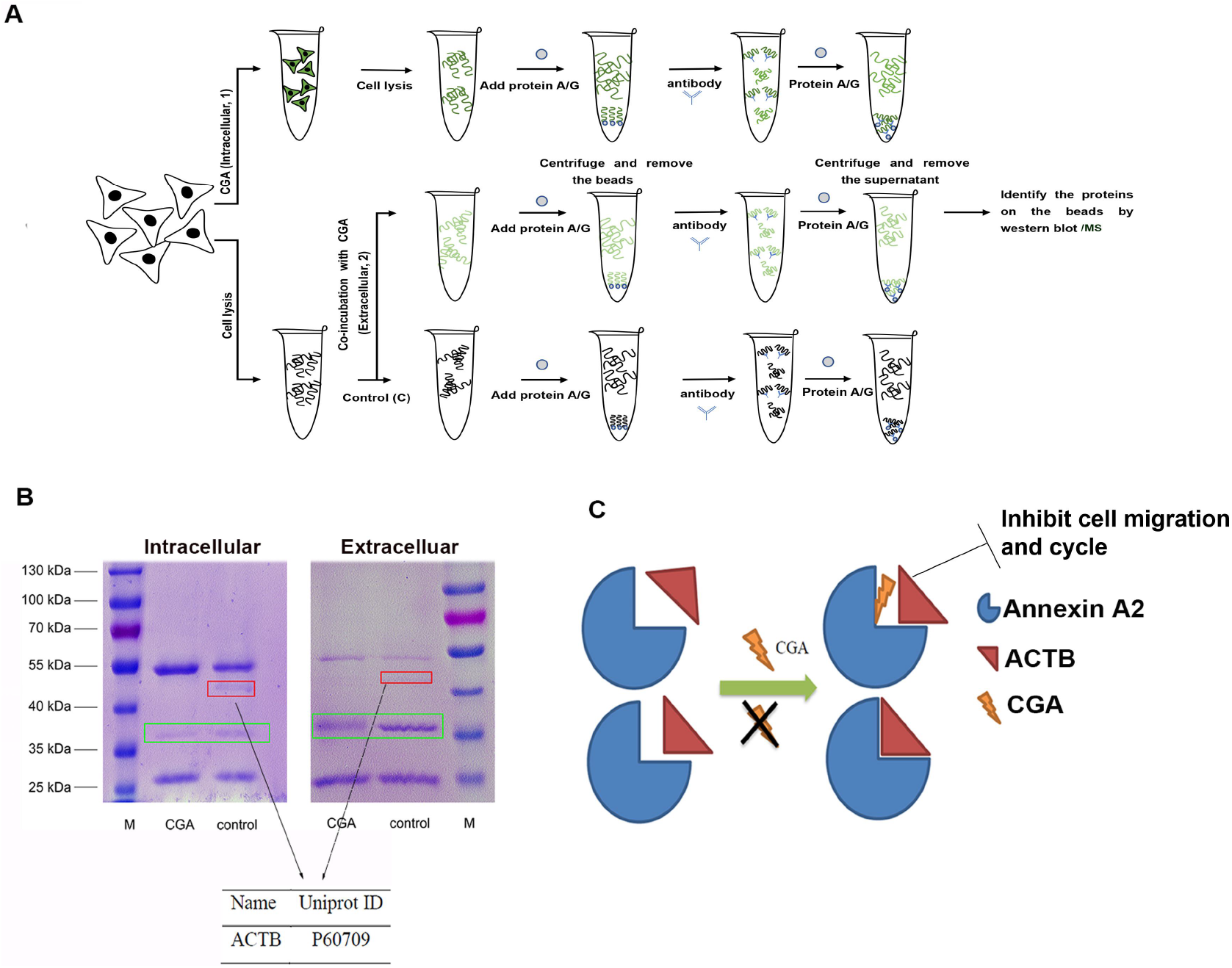
CGA blocked the binding of ANXA2 to actin in A549 cells. **A.** The co-immunoprecipitation was designed carried out for finding the effect of CGA on proteins interact with AXAN2. **B.** After A549 was stimulated with CGA, two protein bands were disappeared. The protein band in the green box is ANXA2 (M: protein maker; control: no CGA added; CGA (Intracellular); CGA (Extracellular)). The protein red box was identified as actin by LC-MS/MS. (C) CGA blocking the binding of ANXA2 to actin may affect cell cycle and migration. MS: Mass spectrometer.

A sandwich ELISA method was developed to detect the ANXA2-actin level in the cell lysate (Figure 8A), the results showed that the level of the complex and the concentration of CGA stimulated cells showed a dose-dependent decline (Figure 8B). The cell scratch test was carried out to test the effect of CGA in inhibiting migration of A549 cells. The healing of scratches had a significant drug-concentration and culture time dependence (Figure 8C). Statistical charts of scratch healing indicated that CGA strongly inhibits the migration of A549 in a concentration and time-dependent manner (Figure 8D). Based on cell viability assay and the cell scratch test, cell cycle test was performed. Compared with control group, the amount of A549 cells treated with CGA increased significantly (from 52.82% to 60.51%) in G0/G1 phase (Figure 8E). It also showed concentration dependent manner and a significant difference compared to the control group. (Figure 8F). These results suggested that the effect of CGA on cell growth is to block cell cycle in G0/G1 phase. The inhibition of migration and cell cycle of A549 cells was confirmed by cell scratch test and cell cycle test. These results correspond to the effect of CGA blocking the binding of ANXA2 to cytoskeleton protein actin.

**Figure 8.**
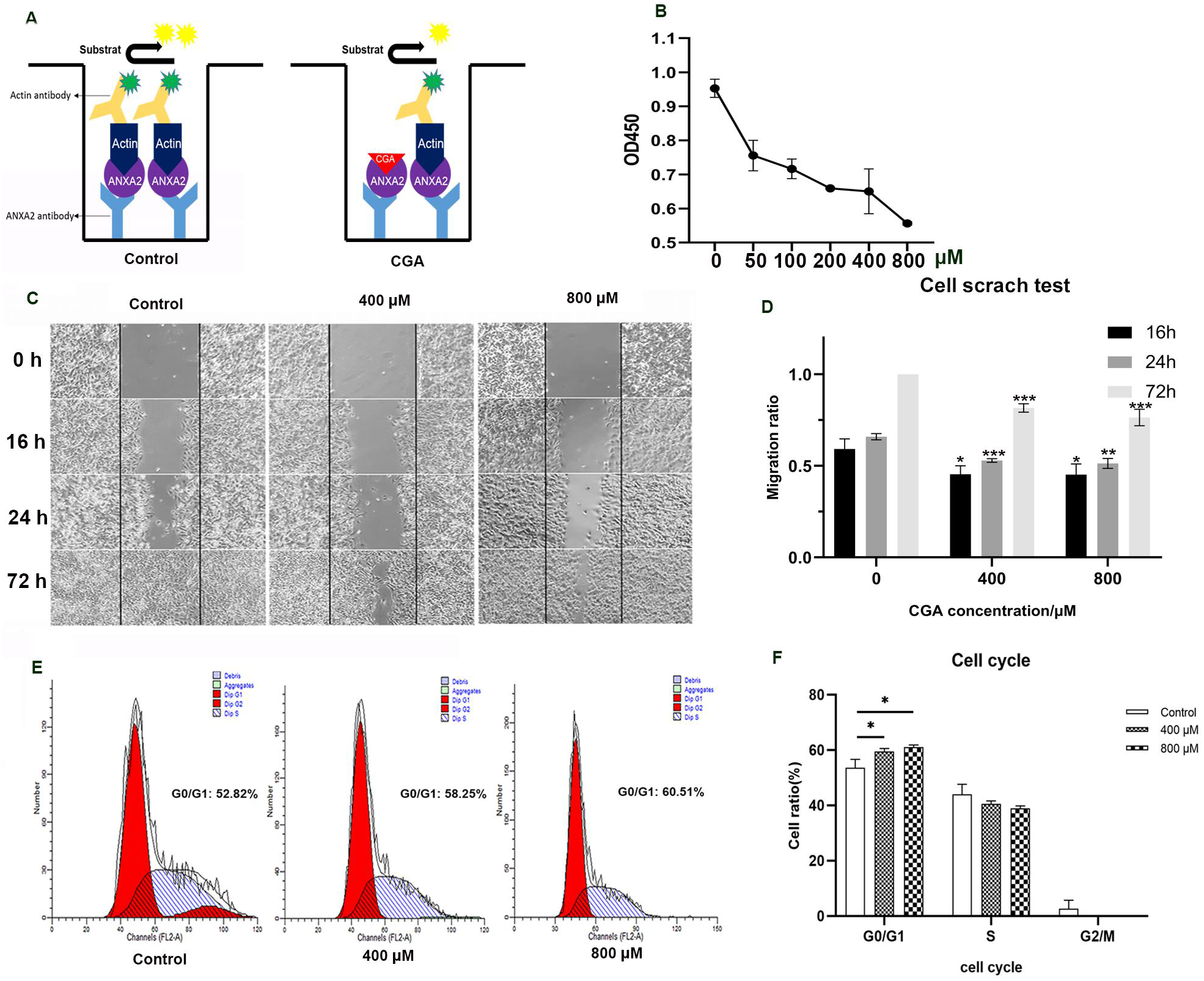
Effects of CGA on cell migration and cell cycle. **A.** The sandwich ELISA method was designed to detect Annexin A2-actin complex. **B.** The level of the complex and the concentration of CGA in stimulated cells is dose-dependent.Each experimental set is representative of n=3 samples and error bars as mean±SD. **C.** CGA inhibited cell migration. When cells were cultured in the medium containing CGA, the healing rate of cell scratch showed a significant concentration dependence of CGA. **D.** Statistical chart of scratch healing ratio of cells cultured with different concentrations of CGA. Repeat three times for each group. At the same time point, different concentrations of CGA treatment group and control group were tested by the t-test statistics. **E.** A549 was cultured for 24 h of 0, 400 and 800 μM CGA, and then the cell cycle was detected by flow cytometry. There was a positive correlation between CGA concentration and G0 / G1 arrest of A549 cells. **F.** Statistical chart of A549 cell cycle was detected by flow cytometry. Repeat three times for each group. Different concentrations of CGA treatment group and control group were tested by the t-test statistics. Data are mean ± SEM. \**P* < 0.05, \*\**P*< 0.01, \*\*\**P* < 0.001

## Discussion

Since screening target proteins of drugs is one of the effective ways to explore their mechanisms, the method of chemical proteomics analysis is widely used. Many effective drug targets were screened by this method, including carnitine palmitoyltransferase 1 (CPT1) as a key target protein of baicalin to improve diet-induced obesity(23) and pyruvate kinase (PyrK) as a target protein of artemisinin to inhibit malaria (24).

In this study, taking advantage of the characteristics of chlorogenic acid (CGA) fluorescence, we proposed the strategy for screening CGA target proteins: using natural proteins with abundant species and complete structure in A549 cells as the natural protein library. ANXA2 was successfully identified as an important target protein for CGA to play its biological functions through the strategy. The identification of drug target proteins from whole cell lysates is a vital work that may be serviceable for the identification and verification of drug targets. The green fluorescence of CGA can be displayed spontaneously as Cy3 like fluorescence (25,26). This greatly avoid false positive binding caused by external modification, increase the credibility of screening results. After ANXA2 was validated as the target of CGA, the MALDI-TOF/TOF result of ANXA2-CGA complex indicates that one ANXA2 molecule binds to one CGA molecule. Based on the structure of the two and the fact that DTT decresed their combination, they are likely to be covalently bound through cysteine by a way similar to Michael addition reaction. ANXA2 is one of the important members of annexin family, which is widely distributed in various eukaryotic cell membranes, cytoplasm and extracellular fluid (27). ANXA2 has a wide range of biological functions mainly involving membrane transport and a series of calmodulin-dependent functional behaviors on membrane surface, including membrane fusion, vesicle transport, cell adhesion, cell proliferation, apoptosis, signal transduction and the formation of some ion channels during exocytosis (26). Overexpression of ANXA2 promotes the survival, growth, migration, angiogenesis and drug resistance of cancer cells (28–31). Based on this target, we try to explain the molecular pharmacology of chlorogenic acid from two aspects: apoptosis and cell cycle change mediated by AXNA2.

ANXA2 can promote the activation of transcription factors NFκB and related factors (32–35), and up-regulates the expression of anti-apoptotic genes, which is conducive to cell survival. ANXA2 is involved in NF-κB signal transduction by combining with p50 subunits and is transported to the nucleus. The presence of ANXA2 enhanced the transcription activity of NF-κB and promoted the downstream anti-apoptotic signal transduction. Ginsenoside, as a binding inhibitor of ANXA2-p50 complex, has also been proved to be effective in inhibiting the proliferation of cancer cells (36). Therefore, CGA is likely to inhibit the development of cancer cells in this way. In our study, CGA, as an inhibitor of ANXA2-p50, also directly disintegrated the ANXA2-p50 complex by binding to ANXA2. Thus, the transcription activity of NF-κB pathway was decreased, in terms of the fact that the transcription and expression of downstream anti-apoptotic genes-cIAP1 and cIAP2 were inhibited. And our animal work also proved that in vivo. After CGA treatment, the size and mass of tumor decreased significantly. Moreover, the anti-apoptotic genes cIAP1 and cIAP2 were also obviously down regulated.

The inhibition of CGA on cancer cells was revealed in the scratch test of A549 migration inhibition and the G0 / G1 phase of cell cycle arrest. Based on the Co-IP results of CGA blocking the binding of ANXA2 to actin, the inhibition of CGA on cancer cells can be partly explained. The binding of ANXA2 to actin is important for the aggregation of actin on cell surface. This interdiction may seriously obstruct cell division and movement(37). As an important cytoskeleton related protein, ANXA2 is highly expressed in a variety of tumor cells(38), including A549 with active proliferation. ANXA2 is related to the proliferation, migration, adhesion and metastasis of tumor cells, and the knockout of ANXA2 can inhibit these processes. Moreover, the specific relationship between the above processes of ANXA2 has not been clearly clarified. The involvement of ANXA2 in these processes may be related to its own interaction with other skeletal proteins (39). In combination with our work, it is also likely to significantly inhibit the migration and cell cycle of tumor cells by blocking the original binding of ANXA2 and actin and other cytoskeleton proteins.

This study may have two potential impacts, on the one hand, it will promote the clinical application of chlorogenic acid. The discovery of protein target will help us to understand the mechanism of drug action. With the help of the expression spectrum of the target in different tumors, the suitable tumor types for the drug can be selected, which will be conducive to accelerate the clinical trials. On the other hand, lung cancer is one of the most common tumors in China. It is of great significance to discover and apply new drugs. The findings of this study will promote AXAN2 as a new tumor target should been paid more attention. The design and development of new drugs based on this target may solve some difficulties in drug development of lung cancer.

It must be point that this study still has some limitations. As a kind of functional molecule, our study on the action of CGA on AXAN2 is still not comprehensive. The influence of chlorogenic acid on the structure of a and whether it will affect other normal functions of AXAN2 are still unknown, although ANXA2 is overexpressed in tumor cells. In animal models, chlorogenic acid has good safety in the treatment and alleviation of tumor(40). What’s more interesting is that in recent years, the safety of chlorogenic acid has been discussed, which is considered to be relatively safe in the treatment of diseases including cancer, inflammation, obesity and diabetes(41,42). Although the results of these studies are encouraging, it should be noted that they are mostly based on the results of animal experiments, and only a small part of them are the data of human related clinical trials. This requires more long-term clinical research to find out the potential danger of chlorogenic acid to human body. This is of great significance for the clinical application of chlorogenic acid.

Our study shows that ANXA2 is an important target protein for chlorogenic acid to play its anti-tumor activity. Through binding to AXNA2, CGA inhibits the activation of NF-κB pathway by interrupting the formation of ANXA2-p50 complex, and finally weakens the transcription and expression of downstream anti-apoptotic genes, such as cIAP1 and cIAP2, to play an anti-tumor role in vitro and in vivo. In addition, CGA blocks the interaction between actin and ANXA2. CGA also arrests cell cycle G0/G1 and obstructs migration of cancer cell. Thus, CGA has the potential to become a significant anti-tumor drug especially in the treatment of lung cancer in the future.

## Experimental procedures

### Cell lines and drugs

As introduced in introduction part, the human on-small cell lung cancer cells (A549) was selected as typical tumor cell model. It was purchased from the American Typical Culture Preservation Center. The A549 cells were cultured in DMEM (HyClone, UT) containing 10% fetal bovine serum (HyClone, UT). Chlorogenic acid (CGA) was purchased from ChemFaces (Wuhan, China). It is dissolved in DMSO (Sigma, MO) at a concentration of 0.28 M and stored at −20 °C.

### Cell viability and proliferation test

The logarithmic growth phase cells with good growth status were inoculated into 96 well plates by inoculating 4×10^3^ cells per well. Each group with 5 repeated pores was cultured. Drug-containing medium (Called DCG) and drug-free medium (Containing DMSO of the same volume as the drug, called DFCG) were added after cell adherence. In addition, cell-free and drug-free control group (Called CFDFG) was set up. After 48 hours of continuous culture, the fresh medium containing cell counting kit-8 assay (CCK-8) (Dojindo, Kumamoto-ken Japan) 10 μl was added to each pore. The cells were cultured for 30 min. OD value of 450 nm was measured by enzyme labeling instrument (The reference wavelength was 600 nm). The cell viability rate was calculated by the formula “cell viability rate = (OD _DCG_ - OD _CFDFG_)/ (OD _DFCG_ - OD _CFDFG_)”. Finally, the cell viability rate curve was drawn.

### Animal work

All the mices were maintained under standard animal housing conditions and free access to food and water. Animal experiments was carried out in accordance with the ARRIVE guidelines and the China Academy of Chinese Medical Sciences Animal Care and Use Committee ethical standards and national guidelines. Six weeks male Balb/c nude mice were obtained from Experimental animal center of Peking University Medical Department (Beijing, China). Nude mice (n=5, each group) were given subcutaneous injection of A549 cells (5×10^6^ cells per mouse) in the left front leg. After injection, the mice were given intraperitoneal injections of 0 mg/kg and 120 mg/kg CGA every day. After 4-weeks treatment with CGA or DMSO, all mice were sacrificed and tumors were removed, weighed, and photographed. Tumor tissues were taken for Western blot.

### Tandem-Mass-Tag (TMT) quantitative proteomics

A549 cells of two groups were cultured with 400 μM CGA and DMSO for 24 h. Cells were collected by 8M urea and the protein concentration was determined by BCA method. 100 μg protein was taken from each group for subsequent experiments. Firstly, the protein samples were treated with 10 mM dithiothreotol (DTT) (Sigma, MO) at 56 °C for 1 h and 55 mM IAA (Sigma, MO) for 45 min. After dilution of urea concentration to 1M, 2.5 μg trypsin (Sigma, MO) was added. Samples were digested overnight by trypsin at 37 °C. The digested samples were desalinated by C18 reversed-phase chromatographic column (Waters, MA) and then vacuum-dried. The samples were dissolved by 100 mM triethylammonium bicarbonate (TEAB) (Sigma, MO) with 100 μl volume. After adding TMT labeling reagents (TMT-126 and TMT-128, Thermo Scientific, MA) dissolved in 41 μl anhydrous acetonitrile to the samples, the samples were well blended. The samples were incubated at room temperature for 1 hour. After that, 8 μl 5% hydroxylamine was added to the samples and then blended. Continue incubating at room temperature for 15 minutes to terminate the reaction. The two groups of samples were fully mixed in equal volume. Finally, the peptides were classified and desalinated by reverse liquid chromatography and analyzed by LC-MS/MS (Thermo Scientific^TM^ Orbitrap Fusion^TM^, MA). The data obtained were analyzed and processed by the bioinformatics tool-string (functional protein association networks: https://string-db.org/), and Cytoscape software (https://cytoscape.org/).

### Chip Spotting and Scanning of CGA

CGA was diluted 100 times and dotted on three-dimensional polymer substrates (CapitalBio, Beijing, China) by chip dotting instrument (CapitalBio, Beijing, China). After that, the green channel (532 nm) was used on a chip scanner (CapitalBio, Beijing, China) to analyze the fluorescence.

### Screening of CGA target proteins by A549 cell lysates and in-gel fluorescence analysis

The cultured A549 cells were lysed with RIPA lysates on ice (Beyotime Biotechnology, Shanghai, China) for 20 minutes. The cell lysates were centrifuged for 20 minutes at 12 000 rpm at 4 °C. The supernatant of cell lysates were incubated with CGA (1 mM) at 37°C for 2 h. After that, it was boiled and SDS-PAGE was carried out. The electrophoresis gel was placed under the fluorescence scanner (GE health care, MA) to complete scanning and imaging.

### In-gel digestion and mass spectrometry

Mass spectrometry and gel digestion were performed as previous. The excised gel pieces were decolorized by 25 mM NH_4_HCO_3_ (Sigma, MO) and 50% acetonitrile (Sigma, MO) and lyophilized in vacuum. To complete the reduction and alkylation, the excised gel pieces were soaked in 25 mM NH_4_HCO_3_ containing 10 mM dithiothreotol for 2 h at 37 °C. After that, gel pieces were placed in 25 mM NH_4_HCO_3_ containing 55 mM iodoacetamide for 45 min in darkness at room temperature. The gel pieces were washed with 25 mM NH_4_HCO_3_ and 50% acetonitrile for 10 min. The excess liquid was absorbed and gel pieces were vacuum freeze-dried. Then gel pieces were treated overnight at 37 °C with 50 mM NH_4_HCO_3_ containing trypsin. The next day, the protein bands were identified by MALDI-TOF/TOF (Bruker, Germany). Finally, the data were analyzed with Mascot bioinformatics database search engine.

### Expression and purification of ANXA2 and p50

The expression and purification method are the same as previously described. Simply, ANXA2 and p50 were overexpressed in *E. coli* BL 21. Then it was lysed by cell ultrasound breaker. Its supernatant was preserved and purified with Ni-NTA resin (Cwbiotech, Beijing, China). The precipitate was dissolved with 8M urea, purified and refolded. The protein concentration was measured by BCA kit (Thermo Scientific, MA). Finally, the protein was identified by MALDI-TOF/TOF.

### SDS-PAGE for testing binding of human recombinant ANXA2 to CGA

The binding of human recombinant ANXA2 to CGA was verified by SDS-PAGE. The recombinant ANXA2 was pretreated as follow and then incubated with CGA (1 mM): (1) The recombinant ANXA2; (2) BSA used as control; (3)The recombinant ANXA2 pretreated at high temperature; (4) The recombinant ANXA2 stored for a long time at ultra-low temperature; (5)The recombinant ANXA2 obtained by renaturation. In addition, ANXA2 was treated with 1, 5, 10, 50 mM dithiothreitol (DTT) at 37 °C for 2 h and then incubated with CGA for 1 h; ANXA2 was pre-incubated with CGA for 1 h and then treated with 10 mM DTT for 2 h. All the above reaction liquids were mixed with protein loading buffer and boiled separately. Then SDS-PAGE was performed. After that, the gel was scanned with a fluorescence scanner or dyed with Coomassie brilliant blue and the results were analyzed.

### Detection of affinity by Isothermal Titration Calorimetry (ITC)

Isothermal drop calorimetry (GE, MA) is a technique used to quantify the interaction of various biological molecules. It can directly measure the heat released or absorbed during the binding process of biological molecules. The recombinant protein annexin A2 and chlorogenic acid were dissolved in the same solvent to eliminate the non-binding exothermic reaction during titration. The concentrations of ANXA2 and CGA were about 12 μM and 200 μM. After that, the isothermal drop calorimeter was set up with suitable parameters (43) to start titration experiment. Finally, the results were analyzed.

### Verify the Binding Ratio of CGA to ANXA2 by MALDI TOF/TOF

The recombinant ANXA2 was incubated with CGA (1 mM) at 37 °C for 2 h. At the same time, recombinant ANXA2 was treated with the same amount of DMSO as control. Subsequently, in order to remove the unbounded CGA and salt, the mixture was eluted and purified by C4 desalination column (Thermo Scientific, MA). Then the molecular weight of the two products was measured by MALDI-TOF/TOF. Finally, the result was analyzed and the binding ratio of CGA to ANXA2 was figured out.

### Immunofluorescence Staining

The clean slide was immersed in concentrated sulfuric acid for 10 minutes. The slide was washed three times with ultra-pure water and sterilized. The cultured A549 cells were digested with trypsin and terminated with DMEM containing 10% fetal bovine serum and dripped onto the slide surface. The slide was carefully placed in a Petri dish and incubated in the incubator for 4 hours. After the cells adhered to the wall, the whole slide was submerged by adding culture medium and cultured overnight.

The slide was carefully washed three times with PBS. Cells were immobilized with 4% paraformaldehyde for 15 minutes. The fixed slide was washed three times with PBS. It was then treated at room temperature with 0.5% Triton X-100 for 20 minutes. After that, it was washed three times with PBS. The excess water on the slide was sucked dry. The slides were sealed with goat serum at room temperature for 30 minutes. Finally, the superfluous goat serum was removed and the slide was stored in a wet box for use.

The cells on slide were incubated with CGA (1 mM) and mouse anti-human ANXA2 monoclonal antibody (Proteintech, Wuhan, China) for 2 hours at 37 °C. Cy5-labeled fluorescent goat anti-mouse antibody (Jackson ImmunoResearch) at 37 °C for 2 hours. Slides were observed under fluorescence microscope (Thermo Fisher Scientific, MA).

### Immunohistochemistry

First, paraffin sections of purchased lung cancer tissues (Best Biotechnology, Xian, China) were dewaxed and hydrated. Then the sections were repaired with 0.01 M sodium citrate buffer at 95 °C. After the slices were washed, 3% hydrogen peroxide solution was added to incubate at room temperature for 20 minutes. Then the slices were washed. The slices were incubated at room temperature for 30 min by appropriate amount of goat serum. After that, the goat serum was thrown off. Mouse anti-human ANXA2 monoclonal antibody (Proteintech, China) was diluted with PBS at 1:50 and dripped onto tissues and incubated overnight at 4 °C in a wet box. Then, the antibody on the slice was thrown off and washed with PBS. After that, tissues were was incubated with CGA (1 mM) at room temperature for 2 h. After that, CGA was washed. Goat anti-mouse Cy5 fluorescent second antibody (Jackson ImmunoResearch) was added to tissues at 1:100 diluted with PBS and incubated at room temperature for 2 h in darkness. Finally, the second antibody on the section was washed. The staining results were observed under fluorescence microscope and photographed for analysis.

### Co-Immunoprecipitation

Protein samples before immunoprecipitation were treated differently. The first protein sample (Extracellular) was derived directly from A549 cell lysates. Then CGA (1 mM) was added to incubate with cell lysates for 1 h, and a control group without CGA added was set up. The 20 μl protein A/G beads (Beyotime Biotechnology, China) were added into the cracking solution and were removed after incubated at room temperature for 1 hour on the blender. After that, the mouse monoclonal antibody of ANXA2(Proteintech, Wuhan, China) of 4 μg was added to the cell lysates and incubated at room temperature for 3 h on the blender. Finally, 20 μl protein A/G beads were added to the cracking solution and incubated at room temperature for 2 h on the blender. After completion, the beads were cleaned three times with pre-cooled PBS. 50 μl protein loading buffer was added immediately and boiled for 5 minutes. SDS-PAGE was performed. The difference between the second protein sample (Intracellular) and the above is that CGA (1 mM) was added in advance to stimulate A549 for 6 hours. The remaining steps are implemented as described earlier. SDS-PAGE was performed.

Same method was used to detect p50-Annnxin A2 complex with mouse monoclonal antibody of ANXA2 (Proteintech, Wuhan, China), and the p50 antibody was detected by Western blot with Rabbit anti-p50 antibody (Proteintech, Wuhan, China).The purified ANXA2 and p50 were mixed equally for 1-2 h on a rotating mixer. Subsequently, CGA (1 mM) was added to the system and mixed on the rotating mixer overnight at 4 °C (at the same time, the drug-free control group was set up). After 20 μl protein A/G agarose beads were incubated with proteins and then removed, mouse anti-human ANXA2 monoclonal antibody of 1 μg was added on a rotary mixer for 3 h. Then the beads were mixed with the above mixing system on a rotating mixer for 4 hours. After that, 20 μl protein A/G agarose beads were washed up by pre-cooled PBS three times. The beads were then boiled for 5 min. Finally, SDS-PAGE was implemented. The results were analyzed.

### Cell Scratch Test

A549 cells in logarithmic growth phase were inoculated into 6-well plate with 1 ×10^6^ cells per hole. When the cells adhere to 90%, the suction head of the 200 μl pipette is used to draw the line vertically in the middle of each hole. Then the six-hole plate was washed three times with PBS to remove the scratched cells. After adding fresh medium containing, 400, 800 μM of CGA and the same amount of DMSO (control group), the cells were cultured in 5% CO_2_ at 37 °C. The initial scratch width and 16, 24, 72 h scratch width of A549 cells were photographed and recorded under a microscope. Scratch repair rate (%) = (initial scratch width - Scratch width at specified time) / initial scratch width × 100%.

### Cell cycle analysis

Cells were inoculated into a six-well plate and were cultured with CGA of 0, 400 and 800 μM for 24h. The cells were then cleaned with pre-cooled PBS and collected. The prepared cells were processed step by step with the cell cycle analysis kit (Beyotime Biotechnology, Shanghai, China). Finally, the cells were detected by flow cytometry (BD Biosciences, NJ).

### Quantitative real-time PCR (QPCR)

Quantitative real-time PCR was implemented as previous (44). After A549 cells were stimulated by 400 μM CGA and DMSO (control) for 24 h respectively, subsequent operation was carried out. Firstly, the total RNA of A549 cells was extracted. Subsequently, the RNA was retranscribed. Finally, quantitative real-time PCR (ABI, CA) was performed. GAPDH, cIAP1 and cIAP2 were detected by q-PCR. The sequence of primers used is shown in Table S1. The data was analyzed by 2^−ΔΔCt^ method.

### Western blotting (WB) and ELISA

A549 cells were stimulated by 400 μM CGA and DMSO (control) for 24 h. Western blotting was carried out. Firstly, the total cell proteins (from A 549 cells or animal tissues in animal work) were extracted and quantified by BCA assay. Then the samples were prepared. After SDS-PAGE was completed, the proteins were transferred to PVDF membrane by electrophoresis. And the PVDF membrane (Millipore, MA) was then sealed in 5% skimmed milk powder for 1 hour at room temperature. After that, the PVDF membrane was immersed in a dilution containing antibodies (ANXA2, NF-κB p105/p50, cIAP1 and cIAP2 specific antibody were purchased from Proteintech (Wuhan, China); actin specific antibody was purchased from Sino biological (Beijing, China) and incubated overnight. After that, the PVDF membrane was incubated with the second antibody (Proteintech, Wuhan, China) at 37 °C for 1 hour after washing 3 times (10 minutes each time) in the shaking bed with PBST (0.2% Tween). Finally, after washing the PVDF membrane (Millipore, MA), the ECL luminescence (APPLYGEN, Beijing, China) detection method is used for imaging analysis.

ELISA method is used for detecting AXAN2-actin complex, 1 ug/mL mouse anti-AXAN2 monoclonal antibody was coated in the 96-well plate (Corning, NY) overnight at 4 °C, then blocked by 2% BSA at 37 °C 1 h. 50 μL A549 Cell lysates(A549 cells were pretreated by 0, 200, 400, 800 μM CGA) and the other A549 Cell lysates(A549 cells were pretreated by DMSO as control) was added into the wells separately at room temperature for 2 h. After washing three times with 0.1% PBST, actin antibody labelled by HRP (Proteintech, Wuhan, China) was added into the well and incubated at room temperature 1 h. After washing three times with 0.1% PBST, TMB reagent (Makewonder Bio, Being, China) added and got the data by ELISA reader (Synergy™, Bio-Tek)

### Computational modelling

The ANXA2 homologous structure model was derived from a part of the Protein Model (PDB ID: 2HYW) in the Protein Structure Database of the National Center for Biotechnology Information (NCBI). The CGA model was also obtained from a structural model (PDB ID: 5GMU) in the same database. The potential binding pockets of ANXA2 were predicted. Then the binding of ANXA2 with CGA was simulated by UCSF Chimera software(45) (www.cgl.ucsf.edu/chimera/). Combining with the experimental data, the possible combination model of CGA-ANXA2 complex was discovered.

### Statistical analysis

Differences between two groups were assessed using two-tailed unpaired t-test. *P* values less than 0.05 were considered significant. \**P*<0.05; \*\**P*<0.01; \*\*\**P*<0.001. Image J (NIH Image) was used to analyze the gray level of the strips and compare the differences in Western Blot.

## Funding and additional information

This study was supported by grants from National Natural Science Foundation of China (81903893), the Fundamental Research Funds for the Central public welfare research institutes (ZZ2018007, ZZ13-YQ-082), Hebei Provincial Department of Science and Technology (19942410G).

## Author Contributions

Lei Wang: Designed and carried out the experiments, data analysis, writing original draft preparation. Hongwu Du: Concept of study, Supervision and revised the manuscript. Peng Chen: Designed and carried out the experiments, Writing-Reviewing and Editing, all authors have read and approved the final manuscript.

## Supporting information

Supporting information related to this article can be found, in the online version.

## Data Availability

All the data is contained within the manuscript and supplemental information.

